# Quantitative Assessment of Morphological Changes in Lipid Droplets and Lipid-Mito Interactions with Aging in Brown Adipose

**DOI:** 10.1101/2023.09.23.559135

**Authors:** Amber Crabtree, Kit Neikirk, Julia Pinette, Aaron Whiteside, Bryanna Shao, Jessica McKenzie, Zer Vue, Larry Vang, Han Le, Mert Dermici, Taseer Ahmad, Trinity Celeste Owens, Ashton Oliver, Faben Zeleke, Heather K. Beasley, Edgar Garza Lopez, Estevão Scudese, Taylor Rodman, Kinuthia Kabugi, Alice Koh, Suzanne Navarro, Jacob Lam, Ben Kirk, Margaret Mungai, Mariya Sweetwyne, Ho-Jin Koh, Elma Zaganjor, Steven M. Damo, Jennifer A. Gaddy, Annet Kirabo, Sandra A. Murray, Anthonya Cooper, Clintoria Williams, Melanie R. McReynolds, Andrea G. Marshall, Antentor Hinton

## Abstract

The physical characteristics of brown adipose tissue (BAT) are defined by the presence of multilocular lipid droplets (LD) within the brown adipocytes and a high abundance of iron-containing mitochondria, which give it its characteristic color. Normal mitochondrial function is, in part, regulated by organelle-to-organelle contacts. Particularly, the contact sites that mediate mitochondria-LD interactions are thought to have various physiological roles, such as the synthesis and metabolism of lipids. Aging is associated with mitochondrial dysfunction, and previous studies show that there are changes in mitochondrial structure and proteins that modulate organelle contact sites. However, how mitochondria-LD interactions change with aging has yet to be fully clarified. Therefore, we sought to define age-related changes in LD morphology and mitochondria-lipid interactions in BAT. We examined the three-dimensional morphology of mitochondria and LDs in young (3-month) and aged (2-year) murine BAT using serial block face-scanning electron microscopy and the Amira program for segmentation, analysis, and quantification. Analysis showed reductions in LD volume, area, and perimeter in aged samples compared to young samples. Additionally, we observed changes in LD appearance and type in aged samples compared to young samples. Notably, we found differences in mitochondrial interactions with LDs, which could implicate that these contacts may be important for energetics in aging. Upon further investigation, we also found changes in mitochondrial and cristae structure for mitochondria interacting with LD lipids. Overall, these data define the nature of LD morphology and organelle-organelle contacts during aging and provide insight into LD contact site changes that interconnect biogerontology and mitochondrial functionality, metabolism, and bioactivity in aged BAT.

Graphical Abstract
Workflow highlighting the process of murine interscapular BAT extraction, processing and imaging serial block facing-scanning electron microscopy and using Amira software for 3D reconstruction of LDs and mitochondrial lipid contact sites to elucidate their structure across aging.

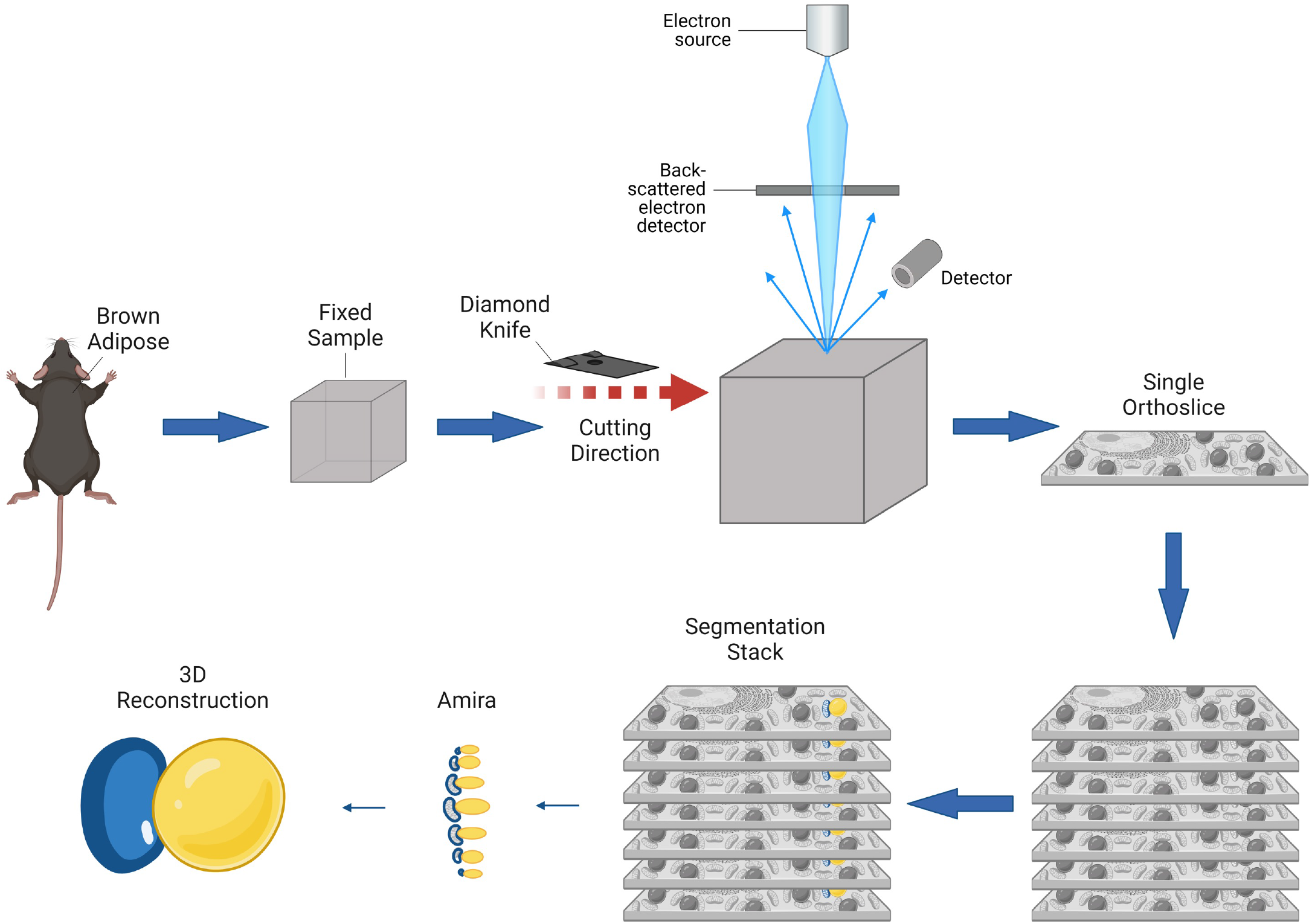

## INTRODUCTION

Mitochondria have numerous indispensable functions, including the maintenance of calcium homeostasis, but they are most often appreciated for their role in energy production (1). Within mitochondrial networks, the function of each mitochondrion is believed to be both tissue and cell-type dependent (2,3). As mitochondrial function is tightly associated with mitochondrial morphology, mitochondrial ultrastructure changes in accordance with the energetic and functional demands of the cell (2,4). Consistent with this specialization, functional and subsequently, morphological, heterogeneity is observed in various tissue and cell types (5). For example, mitochondria typically display a branched phenotype in the transverse orientation and are located between myofibrils in skeletal muscle due to their high energetic needs; in contrast, cardiac mitochondria demonstrate a compact phenotype (6). Mitochondrial heterogeneity is also observed within each cell and this is somewhat dependent on subcellular localization (e.g., position in intermyofibrillar or subsarcolemmal regions) and the dynamic nature of mitochondria (2,3).

Mitochondrial dynamics consist of coordinated cycles of fusion and fission, which provide a means for content mixing, quality control, and the maintenance of healthy mitochondrial networks (7). In brown adipose tissue (BAT), the reorganization of the mitochondrial ultrastructure works cooperatively with the release of free fatty acids to promote heat production via uncoupled respiration (8,9). Immediately following cold exposure or adrenergic stimulation, mitochondrial fission is promoted, which is essential for BAT thermogenesis (9,10). Moreover, recent studies demonstrate the role of mitochondrial architecture in maintaining thermogenic capacity and energy homeostasis in BAT (11–13). During the aging process, a functional decline in BAT is observed and is associated with the onset of metabolic disorders, including type II diabetes (1,2). Given that altered mitochondrial structure, metabolism, and bioenergetics are associated with insulin resistance and type II diabetes (14), the intricate relationship between BAT and mitochondrial structure may offer insight into metabolic pathomechanisms.

Recently, we showed that in murine BAT, aging is associated with a loss of the inner mitochondrial folds, known as cristae, and an increase in mitochondrial volume and matrix (15). Although the high abundance of mitochondria is a notable characteristic of thermogenic adipose tissue, it is not the only defining characteristic. Multilocular LDs, as opposed to the unilocular LDs observed in white adipose tissue, are also a defining feature of thermogenic adipose tissue (16). Studies have shown that the composition and distribution of lipids in BAT can alter mitochondrial function and efficiency (17–19). Additionally, across aging, BAT has been illustrated to accumulate phospholipids, sphingolipids, and dolichols (20), which points toward increased LD formation that may be counteracted by mitochondria-LD-mediated lipolysis (21–23).

LDs are considered dynamic organelles that play a role in various cellular pathways (<u>1)</u>. Mitochondria are known to make contact and associate with LDs and other organelles, allowing for fine-tuning vital processes, such as calcium signaling, lipid synthesis, apoptosis, and the formation of autophagosomes (5,17,23–25). In tissues with a high capacity for oxidative metabolism, mitochondria and LDs show extensive interaction networks. In BAT, mitochondria-LD interactions increase following cold stimulation, presumably to facilitate fatty acid consumption for thermogenesis (16,17,26,27). These interactions are categorized into either dynamic or stable subgroups according to their interaction strength (26,28,29). Dynamic interactions are those that can be disrupted using high-speed centrifugation, which include peridroplet mitochondria, whereas stable interactions remain undisrupted using high-speed centrifugation and are also known as LD-anchored mitochondria (16,17). Although our understanding of mitochondria-LD interactions is evolving, changes in organelle contacts that occur with aging remain incompletely understood.

To better examine how the aging process affects LDs and their associations with mitochondria, we utilized murine BAT and serial block-face scanning electron microscopy (SBF-SEM) combined with 3D reconstruction to quantitatively study these organelle interactions with a relatively high spatial resolution (30–32). As with many techniques in EM quantification, minor errors may arise due to the manual tracing method and the irregularity in mitochondrial shape. In addition, the fine distance between organelle contacts is a source of potential error with many automated systems (33). Thus, to ensure the highest accuracy, we employed manual mitochondria and LD contour tracing using previously validated techniques (34). We hypothesized that mitochondria-lipid interactions and lipid morphology dynamics would change with aging. As the first to study 3D organelle-organelle contacts in adipose tissue across aging, we characterized structural changes that may yield valuable insights into the age-related decline in BAT thermogenic capacity and mitochondrial function.

## METHODS

### Animal Housing and Care

C57BL/6J male mice were housed at 22□ C with *ad libitum* food and water intake and maintained with alternating 12-hr dark/12-hr light cycles. Mice were cared for as previously described by Lam et al. (2021) (35). All animal protocols were approved by the University of Iowa Animal Care and Use Committee.

### SBF-SEM Sample Preparation

All subsequent preparation and analysis were performed by individuals blinded to the experimental conditions of the mice. Interscapular BAT depots were excised from 3-month and 2-year aged mice following euthanasia and fixed in 2% glutaraldehyde in 0.1M cacodylate buffer. A heavy metal protocol was used for processing. Samples were washed with deionized H_2_O (diH_2_O) then immersed in 3% potassium ferrocyanide, followed by 2% osmium tetroxide, at 4□ C for 1-hour each. Samples were immersed in filtered 0.1% thiocarbohydrazide for 20 minutes following diH_2_O wash. Samples were washed again in diH_2_O, then immersed in 2% osmium tetroxide for 30 minutes. Samples were incubated overnight in 1% uranyl acetate at 4□ C prior to incubation in 0.6% lead aspartate for 30 minutes at 60□ C. An ethanol graded series dehydration was then performed prior to samples being infiltrated with epoxy Taab 812 hard resin. Samples were then immersed in fresh resin and polymerized at 60□ C for 36-48 hours. Transmission electron microscopy (TEM) was utilized to identify regions of interest for each of the resultant resin blocks. Samples were trimmed and glued to an SEM stub prior to being placed into a FEI/Thermo Scientific Volumescope 2 SEM. 300-400 serial sections of 0.09 μm thickness were obtained and collected per sample block. The micrograph blocks were then aligned, manually segmented, and 3D-reconstructed using Thermo Scientific Amira Software (Waltham, MA, USA) per previously described (24,34,36).

### Mitochondrial and LD Ultrastructure Calculations and Measurements

Label analyses were performed following manual segmentation of LDs and lipid-associated mitochondria in the regions of interest (ROIs). Each segmented structure was then subject to label analysis using Amira (34). Blinded-3D structural reconstruction from at least three independent experiments was performed to obtain the SBF-SEM murine BAT data. Sequential orthoslices were manually segmented to obtain 300-400 slices for each 3D reconstruction. 50-200 serial sections were then chosen for each 3D reconstruction at approximately equal z-direction intervals. Serial sections were then stacked, aligned, and visualized using Amira software. All videos and volumetric structure quantifications were generated using Amira software. Measurement algorithms that were not already in the system were manually entered. All mitochondria associated with LDs were manually segmented (Figure 2). A total of ∼800 (3-months) and ∼1100 (2-years) mitochondria associated with LDs were collected for quantification from two mice.

**Figure 1:**
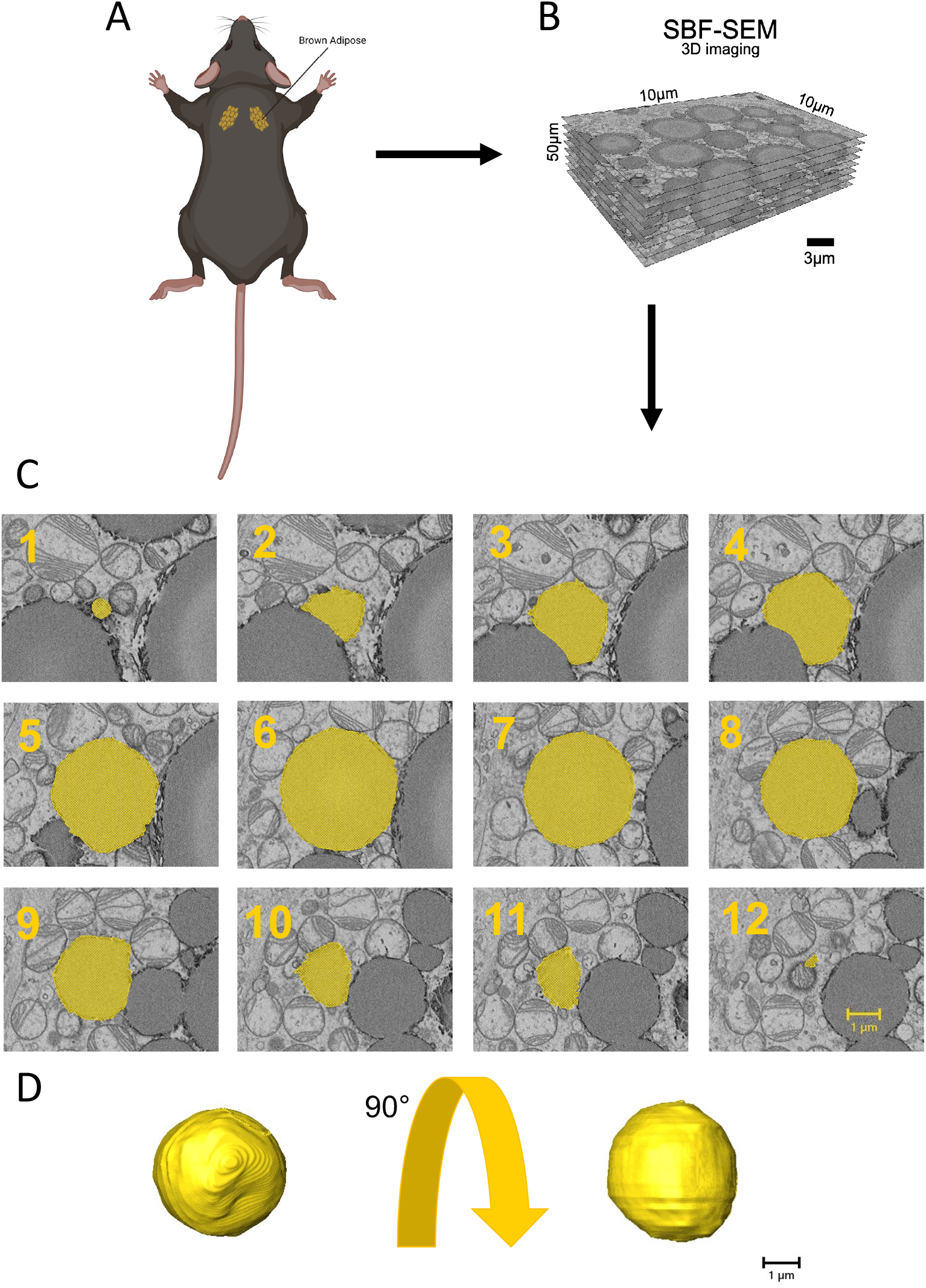
Detailed 3D reconstruction workflow for lipid droplets using serial block face-scanning electron microscopy (SBF-SEM). (A) Interscapular brown adipose tissue (BAT) excised from male C57BL/6J mice. (B) High-resolution 10 μm by 10 μm orthoslices were generated by imaging, aligning, and stacking consecutive ultrathin sections of the sample. (C) SBF-SEM images had lipid droplets manually traced to create representative orthoslices, which provided valuable information on the internal structure of the mitochondria. (D) A detailed 3D reconstruction of lipid droplets was produced based on this workflow, allowing for in-depth analysis of lipid droplet size and complexity.

**Figure 2:**
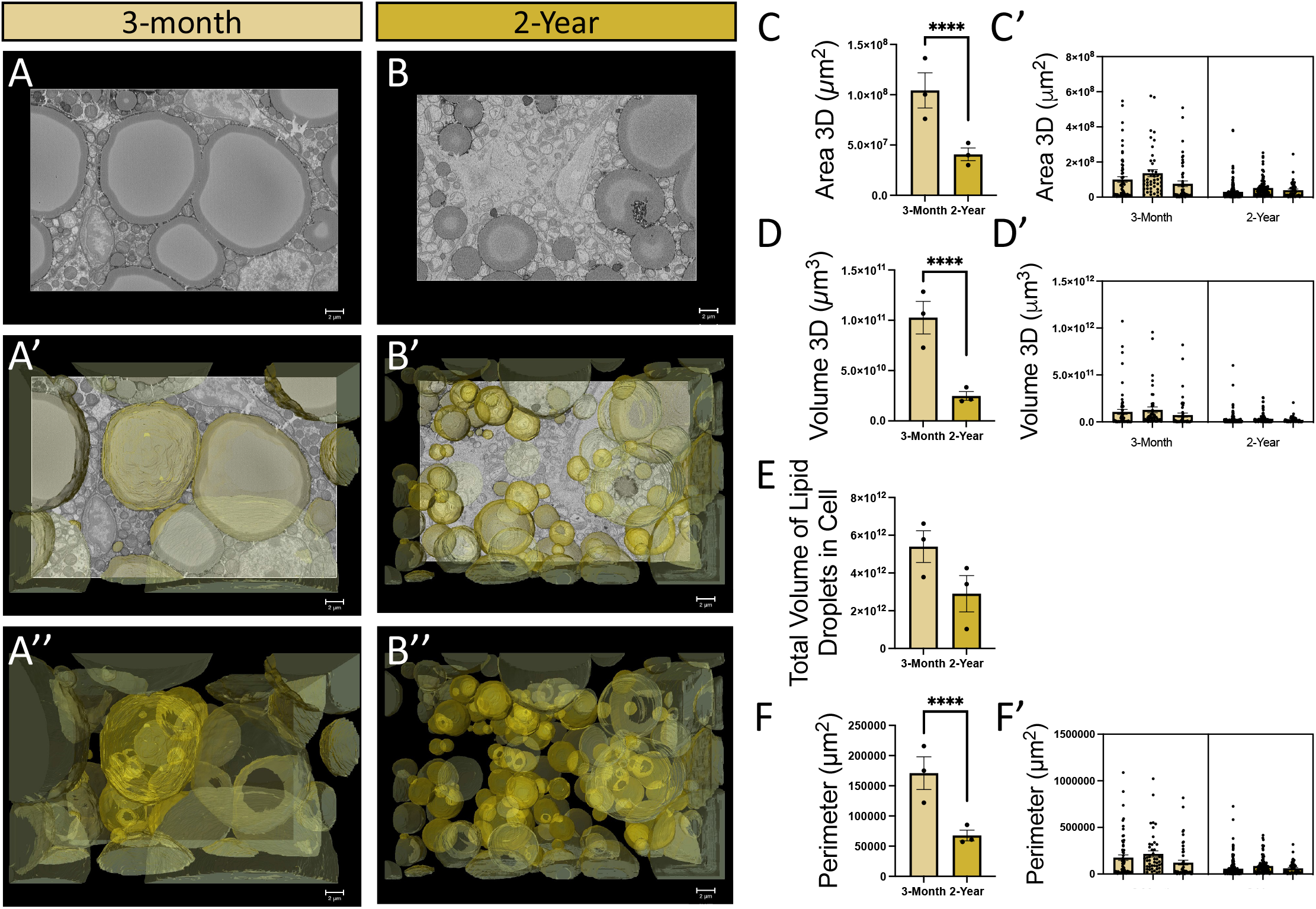
Analysis of mitochondrial morphology changes during the aging process. (A) 3-month-old and (B) 2-year old murine BAT showing: a representative orthoslice; (A’-B’) 3D reconstruction of lipid droplet overlaid on the orthoslice; (A’’-B’’) Isolated 3D reconstruction with orthoslice removed, highlighting the individual mitochondrial structure. (C) Quantitative comparison of 3D lipid droplet area, (D) volume per lipid droplet, (E) cumulative lipid droplet volume, and (F) perimeter between 3-month and 2-year samples to assess morphological differences, revealing age-related changes in size of lipid droplets. Unpaired t-test was used to evaluate changes, with a variable sample number (n approximately 75 per mouse), with each individual mice’s average represented by black dots. (C’, D’, and F’) Display of raw lipid droplet quantifications. Each bar represents one of the three regions of interest surveyed at respective age points while dots represent individual lipid droplets (n = ∼75). Values are displayed for lipid (C’) volume, (D’) area, and (F’) perimeter. In total, 159 lipid droplets across three 3-month male mice and 367 lipid droplets across three 2-year-old male mice were surveyed. The total count of lipid droplets across all mice in respective age cohorts was used for statistical analysis. p < 0.0001 indicated by ****.

### Western Blotting

Tissue from adult (3□months) and aged (2□years) mice were collected, as described above, and lysis occurred with with RIPA lysis buffer (1% NP40, 150□mM NaCl, 25□mM Tris base, 0.5% sodium deoxycholate, 0.1% SDS, 1% phosphatase inhibitor cocktails #2 (Sigma P5726-1ML)/#3 (Sigma P0044-1ML), and one cOmplete protease inhibitor tablet (Sigma 04693159001). After isolating the protein, quantification was performed using a BCA Assay (Thermo Scientific, VLBL00GD2). Equal protein amounts were then separated on 4%–20% Tris-glycine gels (Invitrogen, WXP42012BOX). Following electrophoresis, proteins were transferred to a nitrocellulose membrane (Li-Cor, 926-31092) and incubated overnight at 4°C with primary antibodies: Mic60/mitofilin (Abcam, ab110329). The membrane was incubated for 1 hour at room temperature with secondary antibodies, diluted 1:10,000: donkey anti-mouse IgG (H□+L□) (Invitrogen, A32789) and donkey anti-rabbit IgG (H□+L□) (Invitrogen, A32802). Blots were visualized using the Li-Cor Odyssey CLx infrared imaging system.

### Structure Quantifications and Statistic Analyses

Statistical analyses of all label analysis data and manual measurements were analyzed using a student’s t-test, or the non-parametric equivalent where applicable, in GraphPad Prism (San Diego, California, USA). Individual data points are depicted by dots and all data are considered biological replicates unless otherwise specified. All outliers were used for statistical analyses but may not be displayed for presentation purposes. Graphs are depicted as means with standard error mean (mean ± SEM) indicated as black bars. Statistical significance was defined as p < 0.05 (*), p < 0.01 (**), p < 0.001 (***), and p < 0.0001 (****).

## RESULTS

### Murine BAT Contains Smaller LDs Across Aging with No Changes in Total Cellular Lipid Volume

Cohort age groups were selected as 3 months, which is considered life phase equivalent to 20 years of age in humans, and 2 years, which is considered approximately 69 years of age in humans (37). Following intrascapular BAT tissue extraction, to consider the 3D dynamics of both LDs and mitochondria-LD interactions across aging, SBF-SEM was utilized, as it provides a high spatial resolution (10 μm for the x- and y-planes and 50 μm for the z-plane) while also allowing for large volumes to be surveyed (30). 3D reconstructions were performed using previously described methods (24). For each age cohort (n=3) approximately 150-300 LDs were surveyed, for a total of ∼500 LDs reconstructed (Figure 1A). 3D reconstruction was performed by the manual contour tracing of 50 images of 50-μm block orthogonal (ortho) slices (Figure 1B) across standard transverse intervals (Figure 1C) to enable the reconstruction of pseudocolored LDs (Figure 1D).

Using this workflow of SBF-SEM-based 3D reconstruction, orthoslices (Figure 2A) were overlaid for 3D reconstruction (Figure 2A) of LDs. Based on these representative orthoslices, detailed 3D reconstruction of LDs was rendered (Figure 2A) in both 3-month (young) and 2-year (aged) (Figure 2B-B’’) samples.

Notably, in young samples, we noticed that many individual LDs were too large to fit inside the region of interest, despite the relatively large range of SBF-SEM (10 μm by 10 μm) (30). Although this means that the values for young samples are less reliable, it also supports the notion that aging results in decreased LD volume. This was validated by the significant decrease in LD 3D area (Figure 2C), volume (Figure 2D) and perimeter (Figure 2F) in aged samples. Furthermore, it is unlikely this occurred due to some lipids falling outside of the range of the ROI, as the total cellular volume with lipids showed a marked decrease (Figure 2E). When comparing the LD quantification from each mouse, there was minor inter-group heterogeneity (i.e., differences between samples) but consistent intra-group variability, particularly in the young cohort (Figure 2C’-F’). Together, this indicates an age-dependent loss in LD total size and average size.

Past studies have demonstrated a way to organize mitochondria in a karyotype-like arrangement on the basis of volume (15,38–40). Here, we adapt this method to form a “lipid-otype” organization which allows for the structure and complexity of LDs of various, relatively similar volumes to be compared across the age conditions (Figure 3A). This shows a relatively even distribution of lipid sizes with increased roundness in aged samples. To verify these changes in the roundness of LDs across BAT aging, we calculated lipid sphericity as 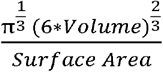). We found that LD sphericity increased with aging in BAT (Figure 3B-B’). Consistent with this, we also found that the complexity, which is inversely related to sphericity, of LD 3D structures decreased with age (Figure 3C-C’). This may indicate that bilayer tension and phospholipid asymmetry are increased, leading to altered LD biogenesis (41).

**Figure 3:**
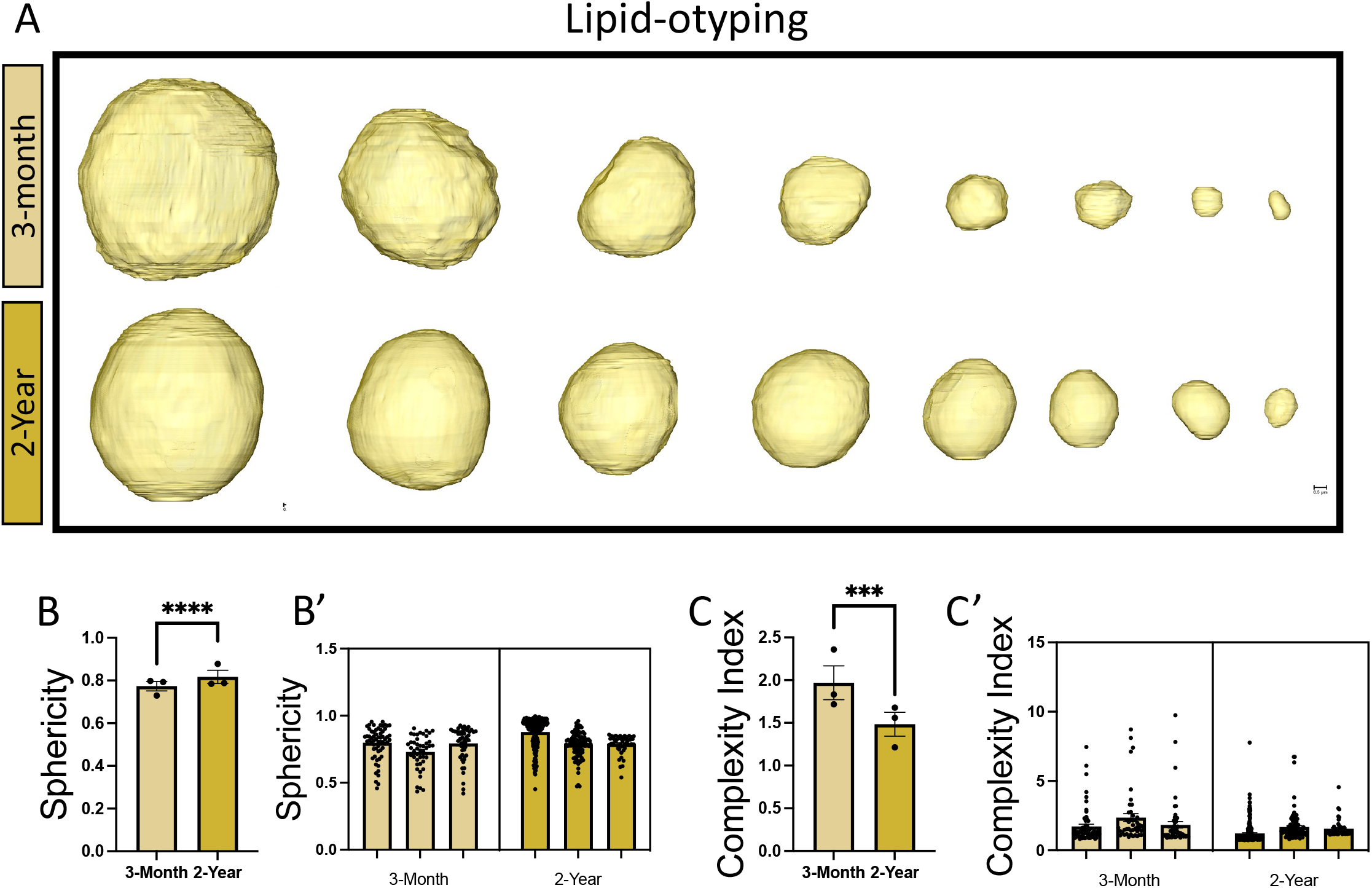
Analysis of age-related changes in lipid complexity. (A) Lipid-otyping for lipid droplet morphology in 3-month and 2-year murine BAT samples based on volume, providing insights into the age-related alterations in structural complexity. These changes in complexity quantitatively comparison of (B) 3D sphericity and (C) complexity index. Unpaired t-test was used to evaluate changes, with a variable sample number, with each individual mice’s average represented by black dots. (B’ and C’) Display of raw lipid droplet quantifications. Each bar represents one of the three regions of interest surveyed at respective age points while dots represent individual lipid droplets (n = ∼75) for (B’) sphericity and (C’) complexity index. In total 159 lipid droplets across three 3-month male mice and 367 lipid droplets across three 2-year-old male mice were surveyed. The total count of lipid droplets across all mice in respective age cohorts was used for statistical analysis. p < 0.001 and p < 0.0001 are indicated by *** and ****, respectively.

Together, these findings show an age-dependent decrease in the size and sphericity of LDs, consistent with changes in morphology. To better understand if the broader cellular environment was concomitantly altered with these morphological changes, we decided to look at possible changes in mitochondrial interactions with LD across aging.

### BAT 3D Reconstruction shows Altered Mito-Lipid Interactions Across Aging

Using the same BAT ROIs that were used for LD quantification, we sought to quantify the mitochondria-lipid interactions in both age groups (Figure 4A-E). Using a similar workflow, BAT was extracted from male mice (Figure 4A) and subjected to SBF-SEM imaging (Figure 4B). LD-associated mitochondria were identified (Figure 4C), and manual contour tracing was done to calculate the contacts between the mitochondria and the LDs (Figure 4D). Notably, we found that mitochondria interacted with up to 4 different LDs simultaneously, owing, in part, to the large volume of LDs in BAT (Figure 4E). Given the high number of interactions, we randomly selected 2 mice from each age cohort (n=2) and surveyed interactions from these samples for approximately 800 total interactions in 3-month samples and 1100 interactions in 2-year samples.

**Figure 4:**
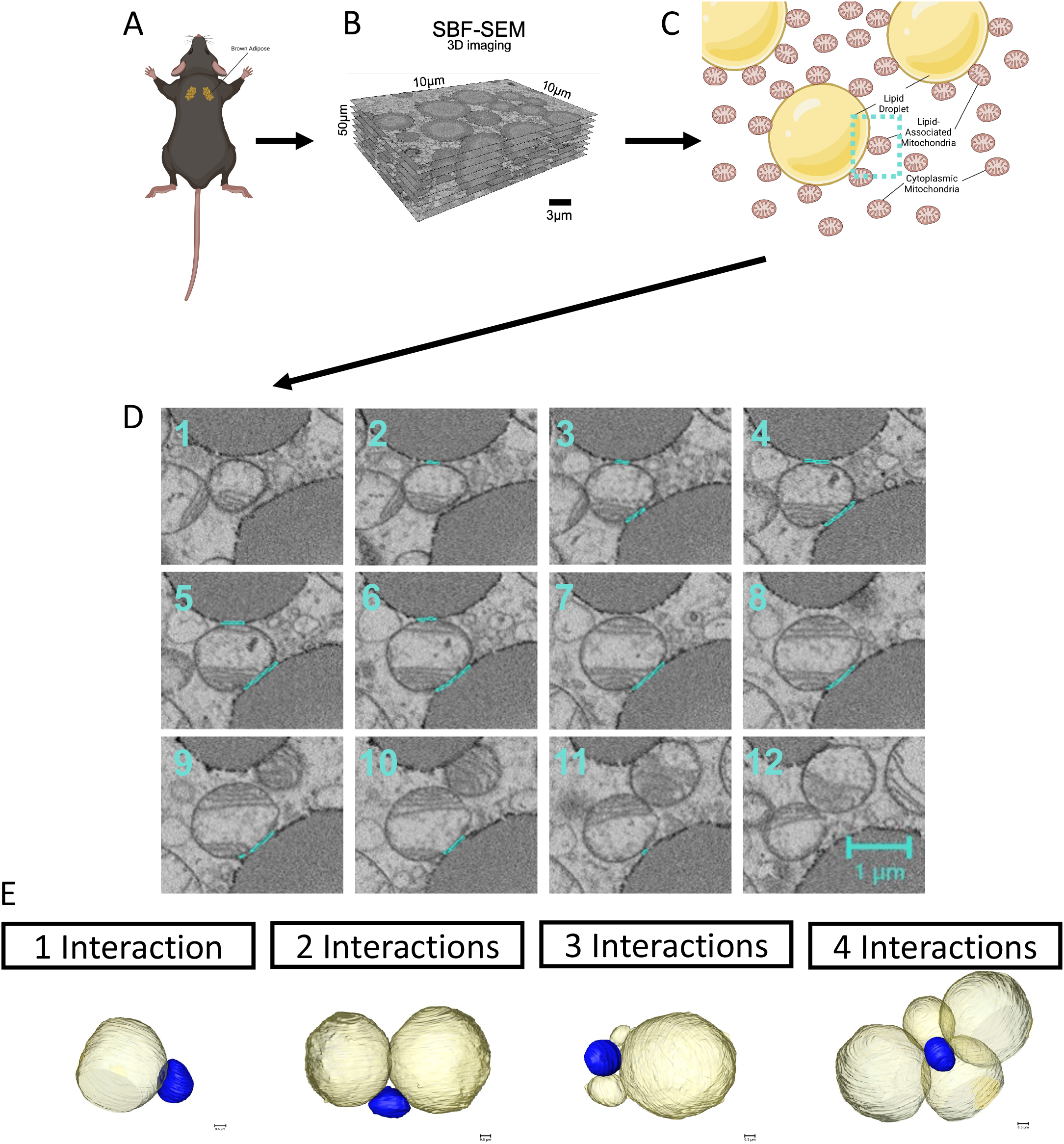
Workflow of serial block face-scanning electron microscopy-based 3D reconstruction for mitochondria-lipid interactions. (A) Brown adipose tissue was excised from 3-month and 2-year aged male C57BL/6J mice. (B) 10 μm by 10 μm orthoslices were overlaid for 3D reconstruction. (C) Lipid-associated mitochondria were identified and (D) manually contour traced, as shown in representative orthoslices from SBF-SEM slices. (E) Diagram of representative images of 1, 2, 3, and 4-lipid droplet interacting mitochondria.

Using this workflow, we reconstructed LDs (Figure 5A’-B’), mitochondria (Figure 5A’’-B’’), and the contact points between the two organelles (Figure 5A’’’-B’’’). We found no statistically significant change in the percent of total mitochondria present interacting with LDs in aged samples (Figure 5C). Interestingly, when disaggregating on the basis of the number of contacts formed, we observed that single interactions were rarer in aged samples, while the proportion of mitochondria interacting with 2, 3, or 4 LD was slightly higher in aged samples (Figure 5D-E). Next, we wanted to see if there were any changes in the mitochondrial and LD surfaces interacting in these aged samples (Figure 5F-L).

**Figure 5:**
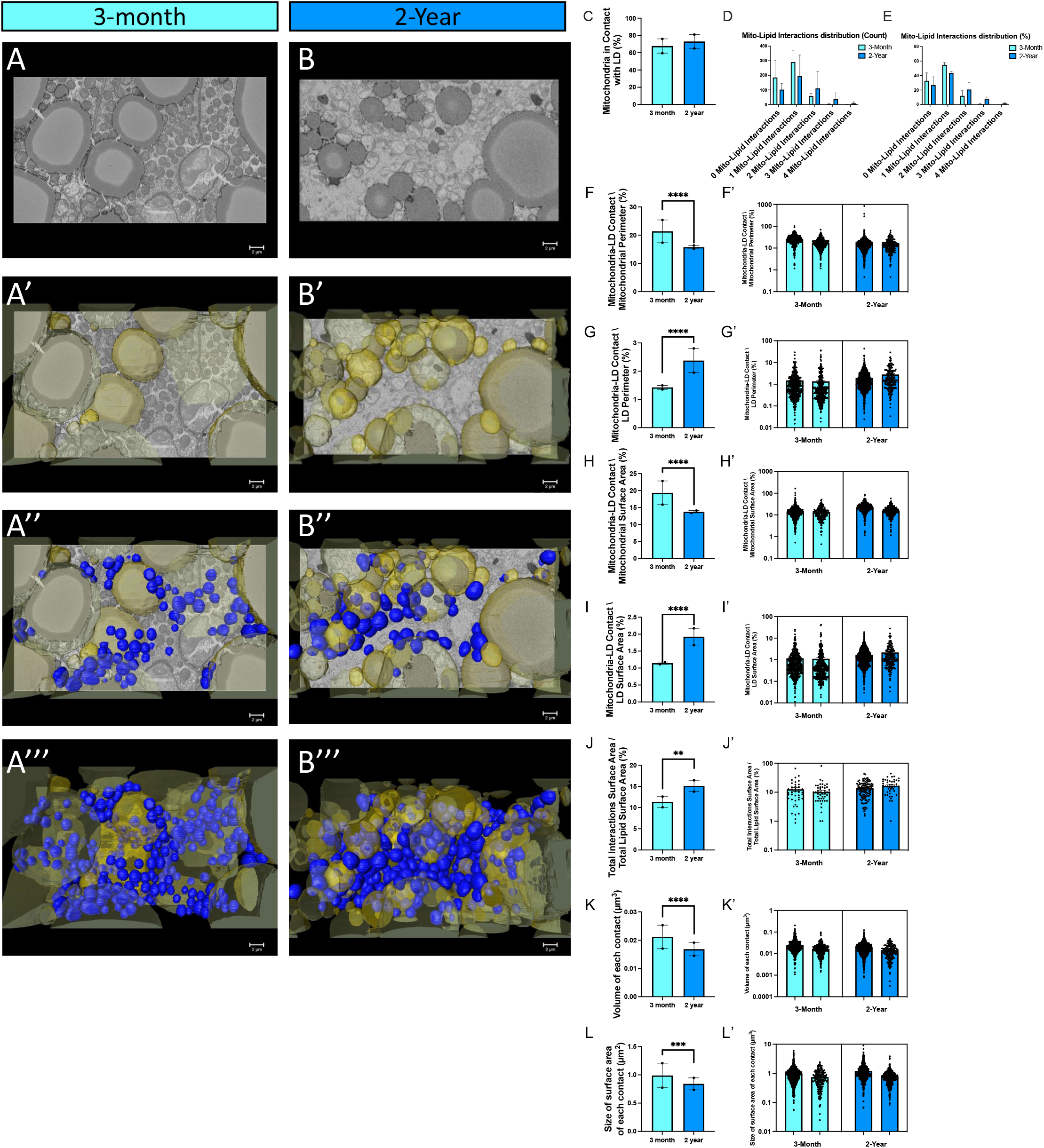
Changes in mitochondria-lipid droplet morphology and frequency across aging. (A-B) Murine BAT representative orthoslice, (A’-B’) 3D reconstruction of lipid droplet (yellow) overlaid on orthoslice, (A’’-B’’) 3D reconstruction of lipid droplet (yellow) and mitochondria (blue) overlaid on orthoslice, and (A’’’-B’’’) 3D reconstruction mitochondria and lipid droplets in (A-A’’’) 3-month and (B-B’’’) 2-year samples. (C-I) Quantification of the lipid droplet-mitochondria contact sites. (C) Comparisons of proportion of lipid-associated mitochondria to cytoplasmic mitochondria. (D) Mitochondrial distribution on the basis of number of lipid interactions formed by count and (E) relative percent. (F) percent contact site perimeter coverage relative to total mitochondrial perimeter, (G) percent contact site perimeter coverage relative to total lipid perimeter, (H) percent contact site surface area coverage relative to total mitochondrial surface area, (I) percent contact site surface area coverage relative to total lipid droplet surface area, (J) percent total contact site surface area coverage relative to total lipid droplet surface area. The average (K) volume of each contact, and (L) surface area of each contact were further elucidated. (F’-L’) Display of heterogeneity in mitochondria-lipid droplet interactions. Each bar represents one of the two regions of interest surveyed at respective age points while dots represent individual contact sites (n = ∼400). Values are displayed for (F’) percent contact site perimeter coverage relative to total mitochondrial perimeter, (G’) percent contact site perimeter coverage relative to total lipid perimeter, (H’) percent contact site surface area coverage relative to total mitochondrial surface area, (I’) percent contact site surface area coverage relative to total lipid droplet surface area, (J’) percent total contact site surface area coverage relative to total lipid droplet surface area, (K’) volume of each contact, and (L’) surface area of contact. Unpaired t-test was used to evaluate changes, with a variable sample number (n approximately 400), with each individual mouse’s average represented by black dots. For “Total Lipid Area in Interactions”, a total of 94 lipid droplet-mitochondria interactions were surveyed across two 3-month male mice, and 163 lipid droplet-mitochondria interactions were surveyed across two 2-year male mice. For all other quantifications of lipid droplet interactions, a total of 839 lipid droplet-mitochondria interactions were surveyed across two 3-month male mice, and 1143 lipid droplet-mitochondria interactions were surveyed across two 2-year male mice. The total count of lipid droplets-mitochondria interactions across all mice in respective age cohorts was used for statistical analysis. p < 0.05, p < 0.01, p < 0.001, and p < 0.0001 are indicated by *, **, ***, and ****, respectively.

Aged samples demonstrated a decrease in the percent of the mitochondrial perimeter and surface area in contact with LDs (Figure 5 F & H). In contrast, the percentage of LD perimeter in each interaction in contact with mitochondria increased significantly with aging, likely due to the smaller size and more abundant presence of LDs in the aged group (Figure 5G). There was a similar age-dependent increase for LD surface area in contacts occupying a higher proportion of total LD volume (Figure 5I). While these quantifications measured only the relative size of single interactions, given many LDs engaged in multiple contacts, we also totaled the surface area of all contacts each LD engaged in and compared it to the overall surface area of the LD (Figure 5J). This mirrors the findings of individual contacts, showing that contacts represented an increased relative to LD surface area, from 10% in young samples to about 15% in aged samples (Figure 5J). Together, this indicates that, with age, a lower percentage of the total surface of mitochondria are dedicated to mitochondria-LD interactions, while the inverse is true for LDs with both individual and cumulative interactions occupying a large proportion of the LD area and perimeter.

Contact sites can be measured by the tightness of their junctions and their impact on modulating the structure of each organelle. Therefore, we examined age-related changes that occur at the contact sites themselves. This led us to discover a significant decrease in the average volume and surface area of each mitochondria-LD contact across aging (Figure 5 K&L). This indicates that the contact sites are decreasing in surface area, with a potential tightening of these contacts resulting in reduced volume. Notably, there was a large sample heterogeneity between the two ROIs surveyed, indicating both intra- and inter-sample variance (Figure 5F’-L’).

### 3D Reconstruction Shows Mitochondria-LD Interactions Do Not Cause Alterations in Mitochondrial Morphology

We questioned if cytosolic mitochondria would differ in morphology, and therefore function, compared to those interacting with LDs (5). Significant contact was defined as mitochondria with >20% of the mitochondria perimeter interacting with LDs (26). It has been reported that subpopulations of mitochondria are functionally distinct from each other (17). While we previously considered all mitochondrial volume in aged BAT samples (15), here, we sought to classify the phenotypes of the mitochondrial subpopulations. Using the same 2 ROIs, manual contour tracing was used for 3D reconstruction in young (Figure 6A) and aged (Figure 6B) BAT. LD-associated and cytoplasmic mitochondria were selected, 3D reconstructed and overlaid on orthoslices (Figure 6A’-6B’) or viewed isolated (Figure 6A’’–6B’’). These interactions were assessed from multiple viewpoints to validate the analysis (Figure 6C-D). Consistent with our previous results (15), mitochondrial length and width both increased with age (Figure 6E-F). Yet, there were generally insignificant differences in mitochondrial width and length due to LD interactions, with intergroup heterogeneity more readily attributed to age-related differences. However, we observed that cytoplasmic mitochondrial width was unchanged with age, while mitochondria LD-associated mitochondria were significantly increased, indicating that age-related changes in mitochondrial size may be more drastic for mitochondrial subpopulations with close proximity to LDs. Together, these results indicate that, while the size of mitochondria-LD interactions changed across aging, the subpopulations of mitochondria may not always dynamically change in response to these alterations in contact sites.

**Figure 6:**
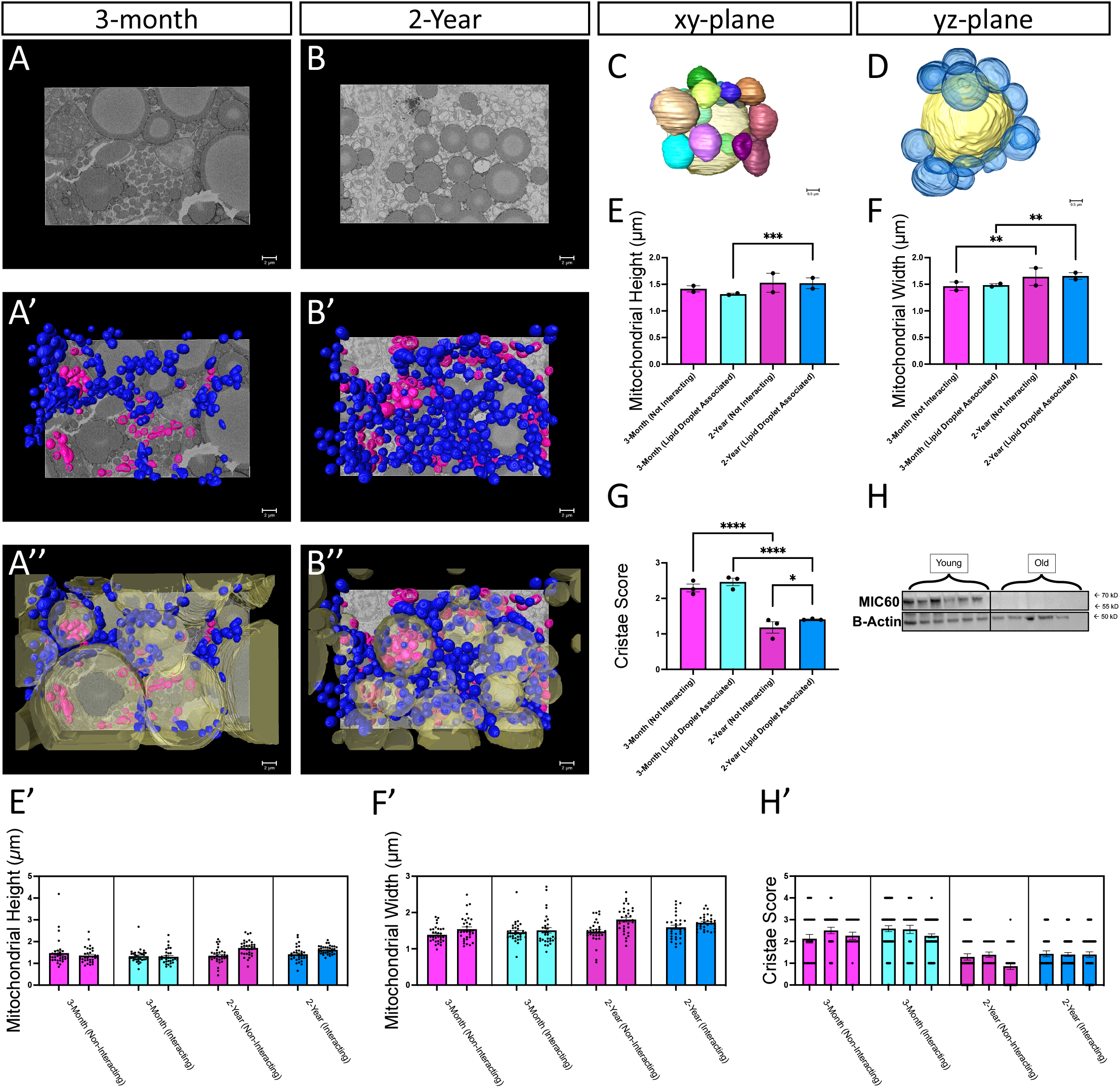
Lipid-Associated Mitochondria Show Distinct Subpopulation Changes in Mitochondrial Architecture. (A) 3-month murine BAT mitochondria viewed from orthoslice, allowing for fine details to be elucidated; (A’) reconstruction of mitochondria and mitochondria associated with lipid droplets (A’’) isolated 3D reconstruction of isolated mitochondria (pink) and lipid-associated mitochondria (blue). (B) 2-year murine BAT mitochondria viewed from orthoslice, allowing for fine details to be elucidated; (B’) reconstruction of mitochondria and mitochondria associated with lipid droplets, (B’’) isolated 3D reconstruction of isolated mitochondria (pink) and lipid-associated mitochondria (blue). (C) Mitochondria associations with lipid droplets viewed from the xy-plane and (D) yz-plane. (E) Comparisons of mitochondrial 3D height and (F) width across aging and on the basis of age and if interacting with lipid droplets. (G) Cristae score, a metric scored from 0-4, of mitochondria compared across aging and interaction status of mitochondria. (H) Western blotting of Mic60 in murine BAT comparing 3-month and 2-year tissue samples. (E’-H’) Variation within mitochondrial and cristae quantifications Each bar represents one of the two or three regions of interest surveyed at respective age points while dots represent individual mitochondria (n = ∼33 for mitochondria; n = ∼50 for cristae, with high variability). Values are displayed for (E’) mitochondrial height, (F’) mitochondrial width, and (G’) cristae score. Kruskal–Wallis one-way analysis of variance with post-hoc pairwise comparison test comparing each mean of groups, was used to evaluate changes, with a variable sample number (n = ∼33 for mitochondria; n = ∼50 for cristae) with each individual mouse’s average represented by black dots. For mitochondrial length and width, in total, 65 mitochondria not interacting with lipid droplets in two 3-month male mice, 65 mitochondria interacted with lipid droplets in two 3-month male mice, 65 mitochondria not interacting with lipid droplets in two 2-year male mice, and 65 mitochondria interacted with lipid droplets in two 2-year male mice were surveyed. For mitochondrial cristae score, in total, 102 mitochondria not interacting with lipid droplets in three 3-month male mice, 173 mitochondria interacted with lipid droplets in three 3-month male mice, 115 mitochondria not interacting with lipid droplets in three 2-year male mice, and 185 mitochondria interacted with lipid droplets in three 2-year male mice were surveyed. The total count of mitochondria across all mice in respective age cohorts were used for statistical analysis. p < 0.05, p < 0.01, p < 0.001, and p < 0.0001 are indicated by *, **, ***, and ****, respectively.

### Mitochondria-LD Interactions may Protect Cristae from Age-related Deterioration

To determine how LD associations may affect mitochondrial bioenergetics, we assessed how cristae structure is altered in aging. To assess cristae abundance and structure, we used a 3D metric of cristae scoring (15,44) assigning a score ranging from 0 to 4 based on the presence and regularity of cristae within the mitochondria. Aging resulted in a significant decrease in cristae quality for both LD-associated and cytoplasmic mitochondrial populations (Figure 6H), consistent with our previous published results (15). While there was no difference in cristae score between LD-associated and cytoplasmic mitochondrial populations at 3 months, comparisons among the 2-year-old cohorts show that LD-associated mitochondrial populations have a significantly higher cristae score. This suggests that LD interactions can ameliorate age-related cristae deterioration, thus providing a potential avenue through which LD interactions restore bioenergetics. Mechanistically, we briefly looked at the MIC60 (also known as mitofillin), a member of the mitochondrial inner membrane complex (MICOS), that regulates cristae structure (42) and morphology (43). We previously reported that levels decrease with aging in cardiac and skeletal muscle tissue (38,39). Therefore, we examined the expression of MIC60 in our BAT aging model and found that levels were decreased (Figure 6H), consistent with our previous studies. This suggests that mitochondria-LD interactions may protect against age-related loss of cristae integrity caused by a decline in the MICOS complex.

## DISCUSSION

In the present study, 3D quantification was used to demonstrate that LDs and the interacting mitochondria exhibit distinct age-related differences in male murine BAT. We found age-dependent losses in size, as well as changes in the distribution and shape of BAT LDs. These findings are contradictory to other reports of increased cytoplasmic LD size in BAT with aging in both male (45) and female (46) mouse models. However, it should be noted that different sub-populations of brown adipocytes exist, with low-thermogenic activity populations exhibiting larger LDs, whereas high-thermogenic brown adipocytes have smaller LDs (47). Additionally, numerous factors could contribute to the variance observed, including, but not limited to, differences in chow diet composition, microbiome, and housing conditions. Moreover, previous studies did not quantify LD size using 3D methods, which likely contributed to the reported differences. Critically, our results indicate that there is increased mitochondria-LD interaction surface area relative to total lipid surface area with aging.

Mechanistically, past lipidomics studies demonstrate that aging causes dysregulated lipid homeostasis, marked by the inhibition of brown adipogenesis concomitantly with the accumulation of certain lipid classes (20). Furthermore, the activity of BAT across the course of life may dictate lipid synthesis, as aging is associated with altered b3 signaling, decreased mitochondrial uncoupling protein 1 (UCP1) expression, as well as altered endocrine functions which may collectively affect lipolysis and thermogenesis in elderly population (48). It may be that the smaller LDs observed are representative of enhanced lipolytic processes within BAT, leading to increased breakdown of triglyceride stored in LDs (48). This increased lipolysis could result in the mobilization of fatty acids from LDs and their subsequent utilization for energy production in thermogenesis. Because mitochondrial lipoylation is particularly decreased in the aging process (49), LDs may have a compensatory role, increasingly utilized for fatty acid metabolism. Therefore smaller LDs may provide a more readily available pool of fatty acids for oxidation compared to larger LDs, enhancing thermogenic capacity. However, as previously reviewed, the complex nature of lipolysis means that smaller LDs are not always indicative of increased lipolysis (50). This is highlighted by conflicting past findings showing decreased markers of lipolysis and blunted thermogenesis in response to cold exposure in aged murine tissue (45). Therefore, defining the cause of smaller LDs in our samples of interest for future mechanistic studies.

Past studies have shown that, with cold treatment, murine BAT increases UCP1, a key protein in thermogenesis that uncouples respiration, as well as cristae biogenesis (47). Furthermore, these alterations in mitochondria were concomitant with increased formation of mitochondria-LD associations (17). In general, UCP1 expression changes resulting in alterations in mitochondria and LD morphology are well elucidated. UCP1 increased expression has also been linked to increase mitochondria-LD droplet formation (51). Caffeine-induced UCP1 activation in beige tissue causes increased mitochondria-LD associations, despite reduced lipid content, alongside increased oxygen consumption rate (51). Past studies have demonstrated that oxidative phosphorylation uncouplers increase mitochondrial volume (52), which we observe here and previously (15). However, UCP1 has reduced activity in BAT across aging (53,54), suggesting there would be reduced mitochondrial volume and LD-mitochondria contact formation, which we did not see (Figures 5–6). It is possible that distinct UCP1-dependent and - independent pathways exist for LD-mitochondria formation. Notably, different subpopulations of adipocytes exist, with low thermogenic activity populations exhibiting larger LDs, while cold treatment causes the dynamic interconversion to high-thermogenic activity adipocytes which have smaller LDs (47). A key hallmark of metabolic syndrome, or greater risk for Type II diabetes caused by a high-fat diet, has been LD accumulation (55). Together, this suggests that UCP1-independent reductions in LDs and the inverse increase in mitochondria-LD contact sites may serve as a compensatory mechanism to avoid metabolic syndrome and increase BAT efficiency.

Notably, these structural changes may also be emblematic of physiological age-related changes. For example, past studies have shown that BAT aging is associated with insulin resistance (54), while Caveolin-1 (Cav-1)-deficiency is associated with insulin resistance in BAT (56). Beyond only increasing mitochondrial respiration, Cav-1 loss resulted in the absence of LD-associated mitochondria (23). This suggests that these structural changes can serve as a pathway in which age-related insulin resistance arises. Furthermore, in aged samples, cristae remain more stable in structure if they have LD interactions (Figure 6H), suggesting that these interactions may be protective. This highlights another mechanism through which LD-associated mitochondria may be relevant for pathogenesis. However, with our focus on offering greater structural insight, the mechanisms driving our results must be studied further.

Interestingly, we also noticed mitochondria-mitochondria interactions, marked by a shortening of conact site distances, occurring between mitochondria interacting with LDs and those not interacting (Figure 5). It is possible this is a tri-organelle contact site, as has previously been reported through ER, mitochondria, and peroxisomes (57). However, the functional relevance of mitochondria-mitochondria contact sites has not been well established. Equally possible, these decreased mitochondrial separation distances may be indicative of post-or pre-mitochondrial dynamic events (58). Similarly, we also noticed many mitochondria interacted with more than one LD. It is possible that bioenergetics differ based on the quantity of LDs interacted with. Notably, we did not consider how mitochondrial quantification changed with the number of interactions. Thus, future studies may explicate if there are significant differences when considering mitochondrial quantification on the basis of LD interaction count or tri-organelle contact.

Past studies have shown that across aging, there is a change in the regulation of metabolites related to energy metabolism, nucleotide metabolism, and vitamin metabolism (53). Notably, *de novo* purine nucleotide synthesis is downregulated across the aging (53). It is possible that the morphology we observe here may contribute to this, as mitochondria are well understood to be involved in *de novo* purine nucleotide synthesis through various intermediates such as amidophosphoribosyl transferase (61). Energy metabolism is also increased across the aging process, while hexokinase 1, a rate-limiting enzyme in glycolysis, is upregulated during aging (53). This suggests that, across aging, mitochondria oxidative phosphorylation may be impaired while LD-mitochondria may serve as a mechanism for fatty acid oxidation as an alternative energy source. Yet, different fatty acids are known to have differential effects on LD-mitochondrial interactions; for example, while palmitic acid overload fuels fatty acid oxidation, oleic acid presented a different phenotype of increased LD-mitochondrial contact sites, yet also increased lipid content, suggesting metabolically inactive mitochondria (62). It remains unclear if these age-related metabolic changes are distinct across peridroplet and cytosolic mitochondria subpopulations. To better explore these age-related alterations, future studies should further examine both age-related changes in mitochondria-LD composition (26,60) and protein levels known to link LDs and mitochondria such as PLN1, PLN5, and MFN2.

As LD-associated mitochondria change in architecture compared to cytoplasmic mitochondria, it is possible that they support the growth of LDs for future triglycerides lipolysis through alterations in bioenergetics. Changes in mitochondrial-LD interactions may affect the efficiency of energy production and thermogenesis, leading to impaired lipid utilization and compromised mitochondrial function in aged BAT (17,26,59). There is a unique balance in BAT with two types of mitochondria and LD interactions: 1) anchored and “stable” LD-mitochondrial contact sites that drive fatty acid oxidation and reduce LD size (16,17,23), and 2) peridroplet mitochondria and “kiss and run” dynamics interactions that have distinct protein levels, engage primarily in pyruvate-derived energy, and expand LDs through the synthesis of TAGs (26,60). Here, while biochemically we could not differentiate between these subpopulations, decreased LD volume with age suggests more anchored mitochondria-LD contact sites. Based on this, the structural differences we observe in LDs may result in an increased fatty acid oxidation (16,17,23). This differs from prior literature showing bound LDs resulting in expanded volume (<u>1)</u> and suggests that the relationship between LDs and mitochondrial interactions, as well as potential roles in fatty acid oxidation, must be further explored.

There are several important limitations of our study that should be noted. First, studies show that there are distinct populations of brown adipocytes with either high or low thermogenic activity. Aside from different expression levels of *Ucp1*, these adipocytes also differ in LD size and mitochondrial content (47). Our study did not account for this heterogeneity in the BAT population, which may contribute to some of the variation observed in organelle morphology. Future studies may consider 3D tissue profiling to see if contact site structure varies across populations of adipocytes or *Ucp1* expression (47). Additionally, while LDs and mitochondria form dynamic contact sites, they can also form tight-bound interactions. These tight-bound cannot be separated using ultracentrifugation and have altered physiological functions centered around LD expansion and oxidative functions (26,27,60,63). However, in contrast to mitochondria-endoplasmic reticulum contact sites, which have clearly defined thickness (25), the distance between dynamic and tight/stable LD-mitochondria interactions remains poorly studied, limiting the ability for differentiation and comparison in 3D reconstruction.

It is also important to note that while the murine model allows for studying BAT across aging, it can also differ from human BAT. Interscapular BAT, as studied here, often does not exist in humans beyond early childhood, while the thermogenic roles of BAT are more essential to small rodents than to humans (64). This makes the roles of human BAT less clear, especially in the context of the age-dependent loss of BAT observed in humans (64). Additionally, only male mice were used in this study. This is important to note as sexual dimorphism has relevance in physiological function, given that BAT from women can display altered expression of certain proteins such as UCP1 (65,66). Past findings have shown that young male mice show a correlation between insulin resistance and adipocyte size, whereas females do not (46). Additionally, those findings suggest that triiodothyronine may maintain UCP1 synthesis in female rats, thus maintaining thermogenic capacity across the aging process in females (46). Furthermore, sex-dependent differences can manifest in distinct aging-dependent responses, such as in puberty, where BAT is more abundant and active in boys (67). While BAT has commonalities across sexes, these model and sex-dependent differences should be considered in the interpretation of these results, and future studies should aim to further elucidate the sex-dependent 3D structure of human mitochondrial organelle contacts in BAT.

Overall, this study defines the age-related changes in LD 3D morphology and mito-lipid interactions, which could collectively contribute to the decline in thermogenic capacity and mitochondrial function in aged BAT. Understanding these changes may provide valuable insights into age-associated metabolic disorders and inform potential therapeutic strategies targeting these dysfunctions. Our studies demonstrate the importance of considering organelle contact sites, rather than focusing solely on mitochondrial morphology when examining structural changes occurring in response to aging.

## Contributions

A.C. performed the experimental procedures, data analysis, interpretation of results, and writing of the manuscript.

K.N. and J.P. contributed equally to this work. They performed the experimental procedures, data analysis, interpretation of results, and writing of the manuscript.

H.L., B.S., A.W. Z.V., L.V., T.C.O., A.O., F.Z., H.K.B., E.G.L., E.S., A.K., S.N., J.L., K.K., B.K., and M.M. were involved in data collection, analysis, and interpretation.

S.M.D., J.A.G., C.W., M.R.M., M.S., H.J.K., S.A.M., A.C., M.D., T.A., T.R., H.L., and A.K.. provided critical feedback, contributed to the writing and revision of the manuscript, and assisted in various aspects of the research.

A.G.M and A.H. Jr. conceptualized the study, provided overall guidance, supervised the research, and revised the manuscript.

## Acknowledgments

Funding by the UNCF/Bristol-Myers Squibb E.E. Just Faculty Fund, BWF Career Awards at the Scientific Interface Award, BWF Ad-hoc Award, NIH Small Research Pilot Subaward to 5R25HL106365-12 from the National Institutes of Health PRIDE Program, DK020593, Vanderbilt Diabetes and Research Training Center for DRTC Alzheimer’s Disease Pilot & Feasibility Program. T-32, number DK007563 entitled Multidisciplinary Training in Molecular Endocrinology to A.C. NSF NRT grant 19-22697 and NSF BPE grant 22-17621 (K. Stassun, PI) to A.C.; T-32, number DK007563 entitled Multidisciplinary Training in Molecular Endocrinology to Z.V. CZI Science Diversity Leadership grant number 2022-253529 from the Chan Zuckerberg Initiative DAF, an advised fund of Silicon Valley Community Foundation (to AHJ). NSF EES2112556, NSF EES1817282, NSF MCB1955975, and CZI Science Diversity Leadership grant number 2022-253614 from the Chan Zuckerberg Initiative DAF, an advised fund of Silicon Valley Community Foundation (to S.D.). This work was further supported by NIH grants K01HL130497 (AK), R01HL147818 (AK), R03HL155041 (AK) and R01HL144941 (AK) NSF grant MCB #2011577I to S.A.M. NIH K01AG062757 to M.T.S. I01 BX005352 from the Dept. of Veterans Affairs Office of Research to J.A.G.

## Videos

**Video 1:** Depiction of z-directional ortho slices used to reconstruct lipid droplets in 3-month murine brown adipose tissue.

**Video 2:** Depiction of z-directional ortho slices used to reconstruct lipid droplets in 2-year murine brown adipose tissue.

**Video 3:** Arc shot of 3D reconstructed lipid droplet overlaid on ortho slice in 3-month murine brown adipose tissue.

**Video 4:** Arc shot of 3D reconstructed lipid droplet overlaid on ortho slice in 2-year murine brown adipose tissue.

**Video 5:** Depiction of z-directional ortho slices used to reconstruct lipid droplet-mitochondrial interactions in 3-month murine brown adipose tissue.

**Video 6:** Depiction of z-directional ortho slices used to reconstruct lipid droplet-mitochondrial interactions in 2-year murine brown adipose tissue.

**Video 7:** Arc shot of 3D reconstructed lipid droplet-mitochondrial interactions overlaid on ortho slice in 3-month murine brown adipose tissue.

**Video 8:** Arc shot of 3D reconstructed lipid droplet-mitochondrial interactions overlaid on ortho slice in 2-year murine brown adipose tissue.

